# Molecular regulation of hypoxia responses by H3K4me3 histone demethylases in *Arabidopsis thaliana*

**DOI:** 10.1101/2025.11.30.691383

**Authors:** Diarmuid S. Ó’Maoiléidigh, Emmanuelle Graciet, Ailbhe J. Brazel

**Affiliations:** Department of Biology, Maynooth University, W23 XY3X, Ireland; Current address: Botany Discipline, School of Natural Sciences, Trinity College Dublin, Dublin, D02 PN40, Ireland

**Keywords:** *Arabidopsis thaliana*, epigenetic regulation, histone demethylases, hypoxia response, RNA-seq, transcriptional regulation

## Abstract

Climate change is increasing precipitation and flooding in many regions, causing major crop losses due to low oxygen (hypoxia) in waterlogged plants. Plant hypoxia responses are known to be regulated through the N-degron pathway, which targets transcriptional regulators for degradation in the presence of oxygen. However, our transcriptomic analyses of *Arabidopsis thaliana* N-degron pathway mutant seedlings show upregulation of only ∼45% of core hypoxia response genes, indicating that additional regulatory mechanisms are involved. In mammals, histone demethylases contribute to hypoxia responses due to their oxygen-dependent activity. Whether histone demethylases regulate plant hypoxia responses has remained unclear. Histone H3 lysine 4 tri-methylation (H3K4me3) promotes gene activation and is removed in plants by demethylases such as JUMONJI14 (JMJ14), JMJ16, and JMJ17. Here, we show that global H3K4me3 levels rise in wild-type *A. thaliana* seedlings exposed to hypoxia. In N-degron mutants, elevated H3K4me3 correlates with activation of hypoxia-responsive genes. We dissected the transcriptomic response to hypoxia and found enrichment of hypoxia response genes and stronger induction of certain core hypoxia response genes in *jmj14/16/17* compared to wild-type seedlings. These data indicate a novel role for H3K4 histone demethylases in regulating molecular hypoxia responses in plants, thereby unfolding new areas for exploration.

**Highlight:** Here we use transcriptomic and protein-based approaches to show a role for H3K4me3 histone demethylases in regulating hypoxia induced transcriptional changes in *Arabidopsis thaliana*.

## Introduction

Oxygen levels are known to fluctuate across multiple scales, from geological timescales to physiological gradients. The great oxidation event caused by the advent of photosynthesis in cyanobacteria >2 billion years ago led to an increase in global atmospheric oxygen levels. During this time, life on earth evolved mechanisms for sensing oxygen levels (Taylor and McElwain, 2010). Today, normal atmospheric oxygen is influenced by both environmental and physiological factors, such as altitude, submergence, respiration rate and tissue thickness (reviewed in (Brazel and Graciet, 2023)). Robust activation of the hypoxia response pathway is critical for plant survival to acute hypoxic stress, such as submergence (Riber *et al*., 2015; Gibbs *et al*., 2018; Weits *et al*., 2021; Fan *et al*., 2023). Hypoxia is also an important developmental signal in plants with roles in germ cell (Tadege and Kuhlemeier, 1997; Kelliher and Walbot, 2012), shoot (Gibbs *et al*., 2018; Weits *et al*., 2019) and root (Shukla *et al*., 2019) development.

Hypoxia responses in plants are regulated by a set of transcription factors (TFs) called the group VII ethylene response factors (ERF-VIIs) (Licausi *et al*., 2011) (Fig. 1A). Under normal oxygen conditions, the ubiquitin/proteasome-dependent N-degron pathway, which includes ARGININE T-RNA PROTEIN TRANSFERASE 1 (ATE1), ATE2, and PROTEOLYSIS 6 (PRT6), sequentially modifies and targets the ERF-VIIs for proteasomal degradation following oxidation of their N-terminal cysteine residue by PLANT CYSTEINE OXIDASE enzymes (PCOs), whose activity is oxygen-dependent (Weits *et al*., 2014). Under hypoxic conditions, ERF-VIIs accumulate and regulate the expression of hypoxia response genes (HRGs), including a set of ∼50 “core” HRGs that are evolutionarily conserved and upregulated in response to hypoxia across multiple plant tissues (Mustroph *et al*., 2009). Although *Arabidopsis thaliana* N-degron pathway mutants constitutively accumulate ERF-VIIs, only 55-70% of the core HRGs are upregulated in these mutants according to published transcriptomic data sets (Gibbs *et al*., 2011). Hence, an unknown mechanism is likely to exist in plants to regulate the remaining 30-45% of these core HRGs.

**Fig 1.**
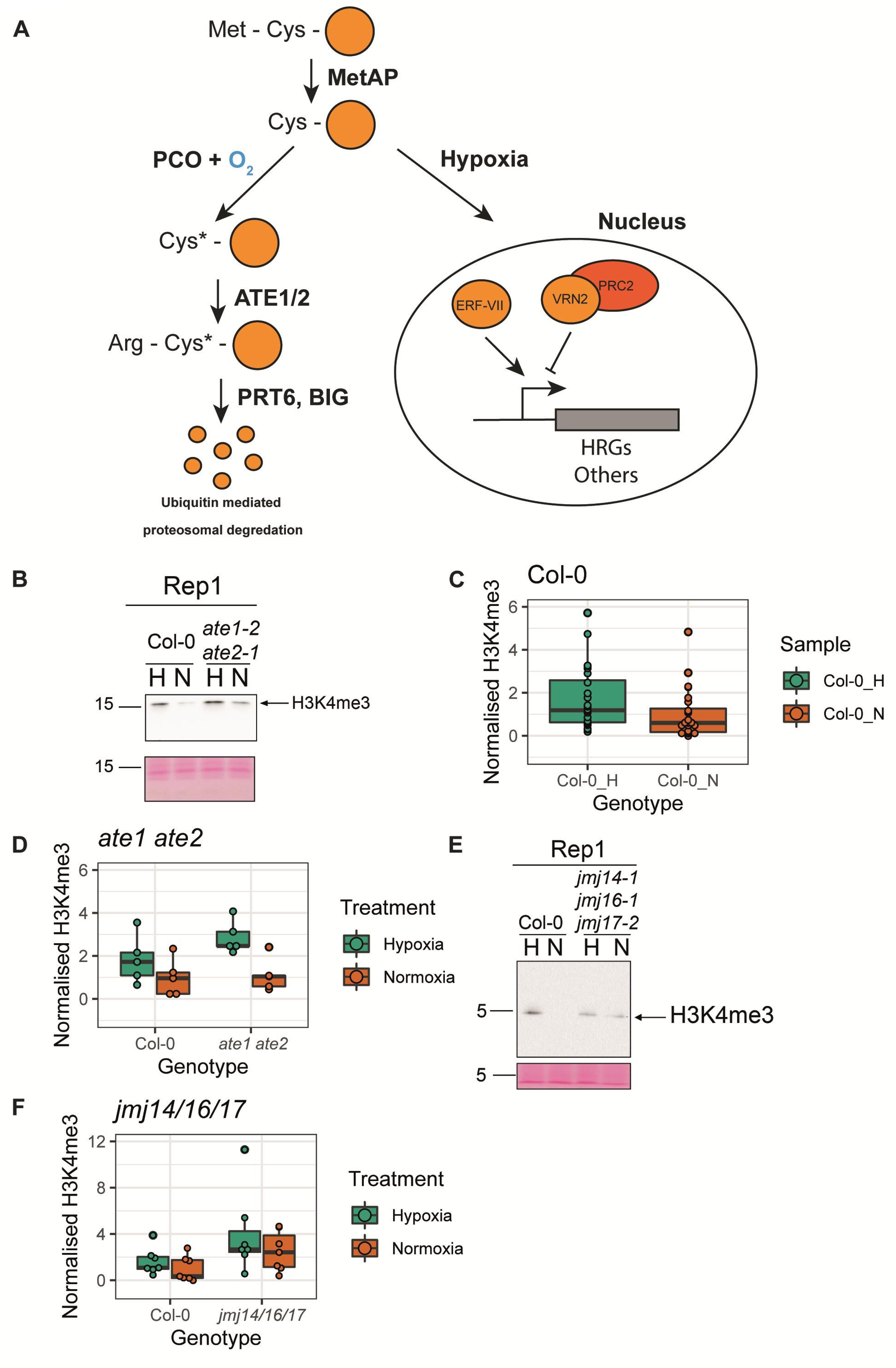
Dynamic changes in global H3K4me3 accumulation in response to hypoxia. (A) Overview of the oxygen-dependent regulation of plant hypoxia responses mediated by the N-degron pathway. METHIONINE AMINOPEPTIDASE (MetAP) enzymes cleave the Met residue from proteins containing N-terminal Met-Cys residues (shown as orange circles) (Giglione et al. 2003). When oxygen is present, PLANT CYSTEINE OXIDASE enzymes (PCO) oxidise the exposed Cys (Cys*). The oxidised Cys is then arginylated by the ARGININE T-RNA PROTEIN TRANSFERASE (ATE) enzymes resulting in an Arg-Cys* N-terminal sequence (White et al. 2017). The N-terminal Arg is then recognised by the E3 ubiquitin ligases PROTEOLYSIS 6 (PRT6) and BIG/DARK OVEREXPRESSION OF CAB1/TRANSPORT INHIBITOR RESPONSE3 (BIG), which polyubiquitinate the protein thus targeting it for proteasomal degradation (Garzón et al. 2007; Zhang et al. 2024). Under low oxygen levels, the PCO enzymes are no longer active and their target proteins escape degradation and accumulate. Targets of the N-degron pathway that accumulate under hypoxia include the ETHYLENE RESPONSIVE FACTOR-VII (ERF-VII) transcription factors, which activate the expression of hypoxia response genes (HRGs) (Licausi et al. 2011). VERNALISATION 2 (VRN2) is a subunit of Polycomb Repressive Complex 2 and another target of the N-degron pathway. VRN2 is involved in the downregulation of target genes via the deposition of histone 3 lysine 27 trimethylation (H3K27me3) through PRC2 (Gibbs et al. 2018). (B) Western blot of H3K4me3 abundance in total protein extracts from 10 d old seedlings treated with hypoxia (0.5% oxygen; H) or normoxia (N) for 8 h, for Col-0 and *ate1-2 ate2-1*. Ponceau S staining of total protein is also shown. (C) Quantification of H3K4me3 in 10 d old Col-0 seedlings treated with hypoxia (0.5% oxygen; H) or normoxia (N) for 8 h normalised to normoxia-treated Col-0 from all Western blots shown in Fig 1 and Fig S1 (n=24). (D) Quantification of H3K4me3 levels in Col-0 and *ate1-2 ate2-1* treated for 8 h normalised to normoxia-treated Col-0 from all Western blots shown in Fig 1B and Fig S1A (n=5). (E) Western blot of H3K4me3 abundance in total protein extracts from 10 d old seedlings treated with hypoxia (0.5% oxygen; H) or normoxia (N) for 8 h for *jmj14-1 jmj16-1 jmj17-2* and Col-0. (F) Quantification of H3K4me3 levels normalised to normoxia-treated 10 d seedlings in Col-0 and *jmj14-1 jmj16-1 jmj17-2* from all Western blots shown in Fig 1E and Fig S1J (n=7). Experiments were performed in independent biological triplicate, each with two biological replicates. Blots for one representative replicate are shown here alongside Ponceau S staining of total protein. Additional replicates can be found in Fig. S1. Details and statistical analyses of replicates can be found in Supplementary Table 2.

Epigenetic regulation in plants plays an important role in the regulation of gene expression and has been found to underpin memory of, and acclimation to, environmental stresses (Iwasaki and Paszkowski, 2014; Kinoshita and Seki, 2014; Gallusci *et al*., 2023). In the context of hypoxia, it has been demonstrated that epigenetic marks, particularly histone acetylation marks, show dynamic changes in their distribution upon hypoxia stress (Lee and Bailey-Serres, 2019). Histone methylation is relevant to hypoxia response as VERNALISATION 2 (VRN2), a component of the polycomb repressive complex 2 (PRC2) that deposits the repressive histone 3 lysine 27 trimethylation (H3K27me3) mark, is a target of the oxygen-dependent N-degron pathway in *A. thaliana* under normoxia (Fig. 1A). In contrast, under acute hypoxia and in developmental hypoxic niches, PCO-dependent oxidation of the N-terminal cysteine of VRN2 no longer occurs, resulting the stabilisation and accumulation of VRN2 (Gibbs *et al*., 2018). When VRN2 is stabilised in hypoxic root tissues, it negatively regulates root growth through the repression of PIF4-regulated transcriptome (Labandera *et al*., 2021; Osborne *et al*., 2025). However, it remains unclear if epigenetic mechanisms are involved in the upregulation of the core HRGs observed in response to hypoxia in plants.

In mammals, the hypoxia inducible factor (HIF) pathway regulates hypoxia responses through the activity of HIF transcription factors. In the presence of oxygen, HIF-α subunits are hydroxylated by the prolyl hydroxylase domain protein 2 (PHD2) and factor inhibiting HIF (FIH) enzymes. This hydroxylation prevents co-activator binding to HIF-α subunits and allows for the binding of the von Hippel Lindau tumour suppressor protein (pVHL) ubiquitin ligase complex which targets the HIF-α subunits for proteasomal degradation (reviewed in Lee 2024). In parallel to PCOs in plants, animals contain an amino-terminal cysteine dioxygenase (ADO) which can oxidise N-terminal cysteines on target proteins, allowing for their recognition and arginylation by ATE1 (Masson *et al*., 2019). HIF1-α was recently identified as a substrate of ATE1, which targets HIF1-α for ubiquitin-mediated proteasomal degradation independently of the pVHL complex (Moorthy *et al*., 2022).

A novel HIF-independent pathway of hypoxia response involving histone demethylases was discovered in mammals (Tausendschön *et al*., 2011; Batie *et al*., 2019; Chakraborty *et al*., 2019). Histone demethylases that contain a Jumonji-C (JMJ) domain catalyse the demethylation of lysine residues on histones and require oxygen as a co-substrate to function, as well as α-ketoglutarate and Fe(II) (Sánchez-Fernández *et al*., 2013). One of the histone demethylases containing a Jumonji-C domain involved in hypoxia response in mammals is KDM5A, which demethylates H3K4me3, an epigenetic mark associated with gene activation (Batie *et al*., 2019). Under hypoxic conditions, KDM5A function is inhibited leading to an accumulation of H3K4me3 and induction of gene expression at a subset of hypoxia-inducible genes (Batie *et al*., 2019). In *A. thaliana*, there are 6 orthologues (JMJ14-19) of mammalian KDM5A/B group histone demethylases (Huang *et al*., 2016). JMJ14-18 have been shown to demethylate marks at H3K4, with JMJ14/16/17 exhibiting demethylation activity for mono-, di- and trimethylated H3K4, JMJ15 showing demethylation activity for trimethylated H3K4 only and JMJ18 showing demethylation activity for both di- and trimethylated H3K4 (Lu *et al*., 2010; Yang *et al*., 2012*a*,*b*; Huang *et al*., 2019; Liu *et al*., 2019).

The JMJ14-19 proteins have been shown to regulate a range of developmental and stress response pathways. JMJ14 (AtPKDM7B) has been shown to regulate flowering time (Lu *et al*., 2010; Ning *et al*., 2015; Zhang *et al*., 2015) and root development (Cattaneo *et al*., 2019), while also being involved in pathogen defence (Li *et al*., 2020) and post-transcriptional gene silencing pathways (Le Masson *et al*., 2012). JMJ14 can co-regulate target genes alongside the NAC transcription factors NAC050/052 (Ning *et al*., 2015; Zhang *et al*., 2015) and can be recruited to target genes by TELOMERE REPEAT BINDING FACTORS (TRBs) (Wang *et al*., 2023), which are also known to recruit PRC2 to target genes (Zhou *et al*., 2018). JMJ15 (AtPKDM7C) is involved in salt stress responses (Shen *et al*., 2014, 2022) and balancing responses to cellular energy and redox state (Zhao *et al*., 2025). JMJ16 (AtPKDM7D) has been shown to repress leaf senescence pathways (Liu *et al*., 2019) and also contributes to the condensation of chromatin during meiosis in male meiocytes, where it can perform cell-type specific demethylation at H3K9 (Wang *et al*., 2020). JMJ17 (AtKDM5) has been shown to play roles in dehydration stress responses (Huang *et al*., 2019), cotyledon greening (Islam *et al*., 2021) and regulation of abscisic acid responses (Huang *et al*., 2019; Wang *et al*., 2021). JMJ18 is primarily expressed in the vascular tissue and has been shown to play a role in flowering time (Yang *et al*., 2012*a*). Despite the demonstrated roles JMJ14-19 histone demethylases play in plant development and stress responses, it remains unknown whether these demethylases contribute to the epigenetic regulation of core HRGs or of other hypoxia response genes.

Here, we show that global levels of H3K4me3 dynamically respond to hypoxia in *A. thaliana* seedlings and present RNA-seq analyses of seedlings with mutations in JMJ14/16/17, alongside seedlings with previously characterized defects in their hypoxia response pathways. Global levels of histone mark H3K4me3 accumulate in response to hypoxia treatment in *A. thaliana* seedlings. We show that genes that are upregulated in response to hypoxia are overrepresented in RNA-seq datasets when comparing *jmj14/16/17* mutant to wild-type seedlings. We also show unique transcriptional signatures of hypoxia response in *jmj14/16/17* mutants, including the higher levels of induction of a selection of core hypoxia response genes. These data suggest a role for H3K4 histone demethylases in regulating hypoxia responses in plants.

## Materials and methods

### Plant growth and materials

*Arabidopsis thaliana* seeds grown on soil were subjected to vapour phase sterilisation using chlorine gas for approximately 3-4 h. Seeds that were grown on MS-Agar were sterilised by washing in 100% ethanol for 5 min followed by 70% ethanol for 7 min. Plants were grown in pots on a compost:vermiculite:perlite (10:6:4) mixture or, for gene expression analysis, plants were grown on an 0.5x MS agar supplemented with 0.5% (w/v) sucrose in Petri dishes. Seeds were stratified at 4°C in total darkness for 2-3 d before transferring to growth rooms at 20 °C, with constant illumination of LED lights.

*A. thaliana* plants of the Columbia-0 (Col-0) accession were used in this study.

The *jmj14-1 jmj16-1 jmj17-2* triple mutant was generated by sequentially crossing plants carrying the T-DNA insertions SALK_135712, SAIL_535_F09 and SALK_037362C. Genotyping PCRs were performed using the genotyping primers listed in Supplementary Table 1. The *ate1-2 ate2-1* and *erf-vii* mutant lines were previously described (Graciet *et al*., 2009; Abbas *et al*., 2015).

### Hypoxia treatment

For hypoxia treatment, seedlings in Petri dishes were removed from the growth room at 9 am and open dishes were placed in a hypoxia chamber (PhO_x Box, Baker Ruskinn) set to 0.5% oxygen and 0% carbon dioxide at 20°C for 8 h in the dark. The corresponding normoxia treated control seedlings were maintained under ambient atmospheric conditions (approximately 21% oxygen and 0.04% carbon dioxide) at 20°C for 8 h in the dark. At collection, plates were removed from the hypoxia chamber and 15-20 seedlings were rapidly collected in liquid nitrogen and stored at -80°C before further processing.

### Protein extraction and Western blotting

Frozen plant tissue was ground to a fine powder using a TissueLyser II (Qiagen). This powder was then resuspended 1:1 w/v in 2X SDS loading buffer (4% SDS, 20% glycerol, 2% 2-mercaptoethanol, 3.2 mM Tris, 0.0001% bromophenol blue). Samples were incubated at 95°C for 10 min and then spun at 18,000 x *g* for 10 min at room temperature, and supernatant was kept for subsequent analysis. Protein concentration was estimated using Amido Black 10B as previously described (Schaffner and Weissmann, 1973; Popov *et al*., 1975) and 20 μg total protein was loaded for each sample on 12% Bis-Acrylamide SDS-PAGE gels. Proteins were transferred to PVDF membranes which were stained using Ponceau S and blocked using 5% (w/v) dried skimmed milk in PBS-T. The primary antibody used in this study for Western blotting was anti-H3K4me3 (1:2000; C15410003, Diagenode). The secondary antibody was anti-rabbit horseradish peroxidase (1:50,000; A0545, Merck). All antibodies were prepared in blocking buffer.

### Quantification of H3K4me3 levels from immunoblots

Relative quantification of H3K4me3 levels was performed using grayscale measurements of protein bands in Photoshop (Adobe, 2022), as previously described (Davarinejad, Hossein, 2015). Ponceau S staining was used as a loading control, whereby Ponceau S bands at either ∼10 kDa or ∼55 kDa were measured for relative quantification. Relative H3K4me3 levels were normalised to the average levels for Col-0 normoxia treated samples for each set of blots. Results of relative quantification are detailed in Supplementary Table 2.

### RNA extraction

Frozen plant tissue was ground to a fine powder using a TissueLyser II (Qiagen). RNA was extracted with a kit (GeneJET Plant RNA Purification Kit, Thermo Scientific) following the manufacturers recommendations. Samples were subsequently treated with DNAseI to remove any residual DNA using a kit (DNA-free DNA Removal Kit, Invitrogen) following the manufacturers recommendations.

### Reverse transcription quantitative PCR

Reverse transcription was performed on RNA to synthesise cDNA using RevertAid H Minus Reverse Transcriptase (Thermo Scientific), RiboLock RNase Inhibitor (Thermo Scientific), dNTPs (Thermo Scientific) and Oligo dT primers (Integrated DNA Technologies) following the manufacturers recommendations. Relative transcript abundance was determined using the Roche LightCycler 480 system and SYBR Green (Roche Applied Sciences). Melting curves were obtained for the reactions, revealing single peak melting curves for all amplification products. The amplification data were analyzed using the second derivative maximum method, and resulting Cp values were converted into relative expression values with the comparative Ct method and using MON1 as a control. Primers that were used for RT-qPCR are listed in Supplementary Table 1.

### RNA-sequencing (RNA-seq)

For RNA-seq analysis, experiments were performed in independent biological triplicate. Quality control, library preparation and paired-ended 150 bp next-generation sequencing with 20 M reads/sample was performed by BGI Genomics (Hong Kong) using the DNBseq sequencing platform. RNA integrity was assessed using an Agilent 2100 Bioanalyzer (Agilent). All RNA samples had RNA integrity (RIN) values between 6.0 and 10 and 28S/18S ≥ 1.

### RNA-seq analysis

RNA-seq analyses were performed using a similar pipeline to that described in (Miricescu *et al*., 2023). The Araport11 release of the *Arabidopsis thaliana* genome annotation was downloaded from The Arabidopsis Information Resource (https://www.arabidopsis.org/) (Cheng *et al*., 2017). Raw RNA-seq reads were aligned to Araport11 using bowtie2 (v2.4.5) (Langmead and Salzberg, 2012). Files were converted from .sam to .bam files and indexed using samtools (v1.15.1) (Danecek *et al*., 2021). Gene abundance was estimated using stringtie (v2.1.7) (Pertea *et al*., 2015) (Supplementary Table 3). Differential gene expression analysis was performed using the Bioconductor package DeSeq2 (Love *et al*., 2014) in R version 4.1.2 (R Core Team, 2021) using a design in which the factors Genotype and Treatment were combined into a single factor to model multiple condition effects. DeSeq2’s median of ratios method was used to normalise read count values. The results of multiple comparisons were extracted and differentially expressed genes (DEGs) were defined by adjusted *p*-value ≤ 0.05 and |log_2_(fold change)| ≥1 (Supplementary Table 4-7). Principal component analysis (PCA) was performed using pcaExplorer (v2.20.2) (Marini and Binder, 2019) in RStudio (version 2022.2.0.443). Gene ontology (GO) analysis was performed using ShinyGO (v0.82) (Ge *et al*., 2020). Venn diagrams were generated using InteractiVenn (Heberle *et al*., 2015). Plots were generated using ggplot2 (v3.3.6) (Wickham, 2016) and modified for style in Adobe Illustrator 2022.

### K-means clustering

The means of normalised read counts were filtered for k-means clustering as follows. Deseq2 DEG analysis was performed, and results for the 16 comparisons shown in Supplementary Table 8 were extracted. A list of 6405 DEGs (filtered by adjusted *p*-value ≤ 0.05 and |log_2_(fold change)| ≥1) was generated by combining DEGs from hypoxia to normoxia samples from the same genotype, DEGs from each normoxia to every other normoxia sample, and DEGs from each hypoxia to every other hypoxia sample. Next, the DEGs with a mean of <10 normalised reads were removed leaving 5926 genes for clustering analysis. Clustering analysis was performed on normalised read counts from the Col-0 and *jmj14-1 jmj16-1 jmj17-2* samples using the k-means function in R (v4.1.2) with the arguments centers = 12, nstart = 1000, iter.max = 300, and algorithm = “Lloyd”. The resulting gene cluster assignment is annotated in Supplementary Table 5-7. Clustering heatmaps were generated using ComplexHeatmap (v2.10.0) (Gu *et al*., 2016) in R.

### Statistical analysis of protein or gene expression levels

Statistical analyses were performed in R on normalised protein levels, normalised read counts or relative expression values for individual genes. A Levene test was performed and confirmed the homogeneity of variances in all the data-sets analysed. A two-way ANOVA was performed using the parameter “Genotype x Treatment”. Tukey Honest Significant Differences (HSD) tests were then performed to test for significant differences between groups. The results of these tests are presented in Supplementary Table 2 and Supplementary Table 9.

## Results

### Histone mark H3K4me3 dynamics in wild type seedlings and seedlings with perturbed hypoxia responses

Previous studies detected a mild decrease in H3K4me3 levels at gene bodies in response to 2 h hypoxia using chromatin immunoprecipitation combined with next-generation sequencing (Lee and Bailey-Serres, 2019). However, we could reproducibly observe an accumulation of H3K4me3 deposition in wild-type *A. thaliana* 10 d old seedlings exposed to hypoxia (0.5% oxygen) for 8 h in comparison to wild-type seedlings kept under normal atmospheric conditions (normoxia; ∼21% oxygen) (Fig. 1B). Levels of H3K4me3 increased by an average of 70% in hypoxia-treated wild-type seedlings (accession Col-0; *p*-value = 3.79e-05; Fig. 1C, Supplementary Table 2). This finding is consistent with the putative oxygen dependent activity of the histone demethylases.

To further explore a potential role of dynamic H3K4me3 in the regulation of the transcriptional response programme, we tested global H3K4me3 accumulation in response to hypoxia in mutant genotypes where the hypoxic response was perturbed. We examined *ate1-2 ate2-1* double mutant seedlings, in which the hypoxia response pathway is constitutively active through disruption of the N-degron pathway and constitutive stabilisation of the ERF-VIIs (Gibbs *et al*., 2011; Licausi *et al*., 2011). Hypoxia treatment reproducibly led to an increase in the levels of H3K4me3 in *ate1-2 ate2-1* (two-way ANOVA, *p*-value = 0.0052; Fig. 1B, Fig. 1D, Fig. S1A, Supplementary Table 2). Median H3K4me3 levels were approximately 2.4-fold higher in *ate1-2 ate2-1* seedlings treated with hypoxia *versus* normoxia (p-value = 0.0318; Fig. 1D, Supplementary Table 2), while H3K4me3 levels were induced by just 1.8-fold in Col-0 seedlings treated with hypoxia (Fig. 1D). These data suggest a positive correlation between the activation of the hypoxia response pathway and global H3K4me3 accumulation, and that constitutive accumulation of ERF-VIIs in the *ate1-2 ate2-1* mutant background contributes to a further increase in global H3K4me3 in response to hypoxia.

We next tested whether global H3K4me3 changed in response to hypoxia in seedlings in which hypoxia response is disrupted through T-DNA insertions in the five *ERF-VII* genes (quintuple mutant noted *erf-vii*) (Abbas *et al*., 2015). We observed variation between biological replicates with the *erf-vii* mutant, as global levels of H3K4me3 accumulated in *erf-vii* seedlings treated with hypoxia in two biological replicates, while in another replicate the opposite result was observed (Fig. S1B). Quantification of the signal indicates that the increase in H3K4me3 upon hypoxia is dampened in *erf-vii* seedlings (Fig. S1C, Supplementary Table 2). Hence, the data suggest that the ERF-VII transcription factors may be involved in the accumulation of global H3K4me3 under hypoxia, but may not be required for this process.

### Contribution of H3K4me3 histone demethylase genes to H3K4me3 dynamics in response to hypoxia

Histone demethylases belonging to the KDM5A/B group have been shown to be involved in regulating hypoxia responses in mammalian cells (Tausendschön *et al*., 2011; Batie *et al*., 2019; Chakraborty *et al*., 2019). As mentioned above, six different loci encode KDM5A/B group histone demethylases in *A. thaliana*. JMJ19 (AtPKDM7A) lacks conserved residues and domain required for demethylase enzymatic activity (Lu *et al*., 2008), while H3K4me3 demethylase activity has previously been demonstrated for JMJ14-18 (Lu *et al*., 2010; Yang *et al*., 2012*a*,*b*; Huang *et al*., 2019; Liu *et al*., 2019). The expression of *JMJ15* is very low in normoxic rosette leaves (Fig. S2A), and *JMJ18* expression is restricted to companion cells (Yang *et al*., 2012*a*). In contrast, *JMJ14*, *JMJ16* and *JMJ17* are broadly expressed at a relatively high level (Fig. S2) and the corresponding wild-type proteins have been shown to have demethylase activity (Lu *et al*., 2010; Huang *et al*., 2019; Liu *et al*., 2019). Therefore, to test a potential role of KDM5A/B-related proteins in hypoxia in plants, we focused on testing global H3K4me3 levels in mutants with T-DNA insertions in *JMJ14*, *JMJ16*, or *JMJ17*.

The H3K4me3 levels were similar in *jmj16-1* seedlings and in the wild type in normoxic conditions, and the average fold induction of H3K4me3 in response to hypoxia was also similar in the two genotypes (Fig. S1D-E, Supplementary Table 2). Median levels of H3K4me3 were moderately higher in *jmj17-2* seedlings compared to Col-0 in both normoxic and hypoxic conditions (Fig. S1F-G). They also increased in hypoxia-treated *jmj17-2* compared to the same genotype kept under normoxic conditions. In contrast, median H3K4me3 levels were approximately 3.6 times higher in *jmj14-1* seedlings in comparison to Col-0 under normoxic conditions (Fig. S1H-I, Supplementary Table 2). However, we did not observe a robust induction of H3K4me3 deposition in hypoxia treated *jmj14-1* seedlings compared to Col-0 seedlings (Fig. S1H-I, Supplementary Table 2). This suggests that *JMJ14* plays a role in the global hypoxia-induced accumulation of H3K4me3 at this stage of development.

To investigate the redundancy between these genes, we generated a *jmj14-1 jmj16-1 jmj17-2* triple mutant line and tested global accumulation of H3K4me3 in response to hypoxia. As observed for *jmj14-1* seedlings, H3K4me3 levels were elevated in normoxia in comparison to Col-0 and did not respond robustly to hypoxia treatment (Fig. 1E-F, S1J). Overall, the behaviour of the triple mutant in comparison to wild-type seedlings was shifted towards increased H3K4me3 deposition (*p*-value = 0.0238; Fig. 1F, Supplementary Table 2). Together these data indicate that *JMJ14* plays a major role in preventing H3K4me3 accumulation under normoxia, whereas *JMJ16* and *JMJ17* may play more minor roles.

### Transcriptional signatures of *jmj14-1 jmj16-1 jmj17-2* seedlings in response to hypoxia

To understand how perturbed H3K4me3 deposition under normoxia and hypoxia influences gene expression, we used RNA-seq to monitor genome-wide gene expression differences of 10 d-old *jmj14-1 jmj16-1 jmj17-2* and Col-0 whole seedlings subjected to 8 h hypoxia (0.5% oxygen, H) or normoxia. A PCA showed that samples separated based on treatment (PC1: 85.73% variance) and genotype (PC2: 9.26% variance), suggesting differences in the way these genotypes respond to hypoxia treatment on a transcriptional level (Fig. 2A).

**Fig. 2.**
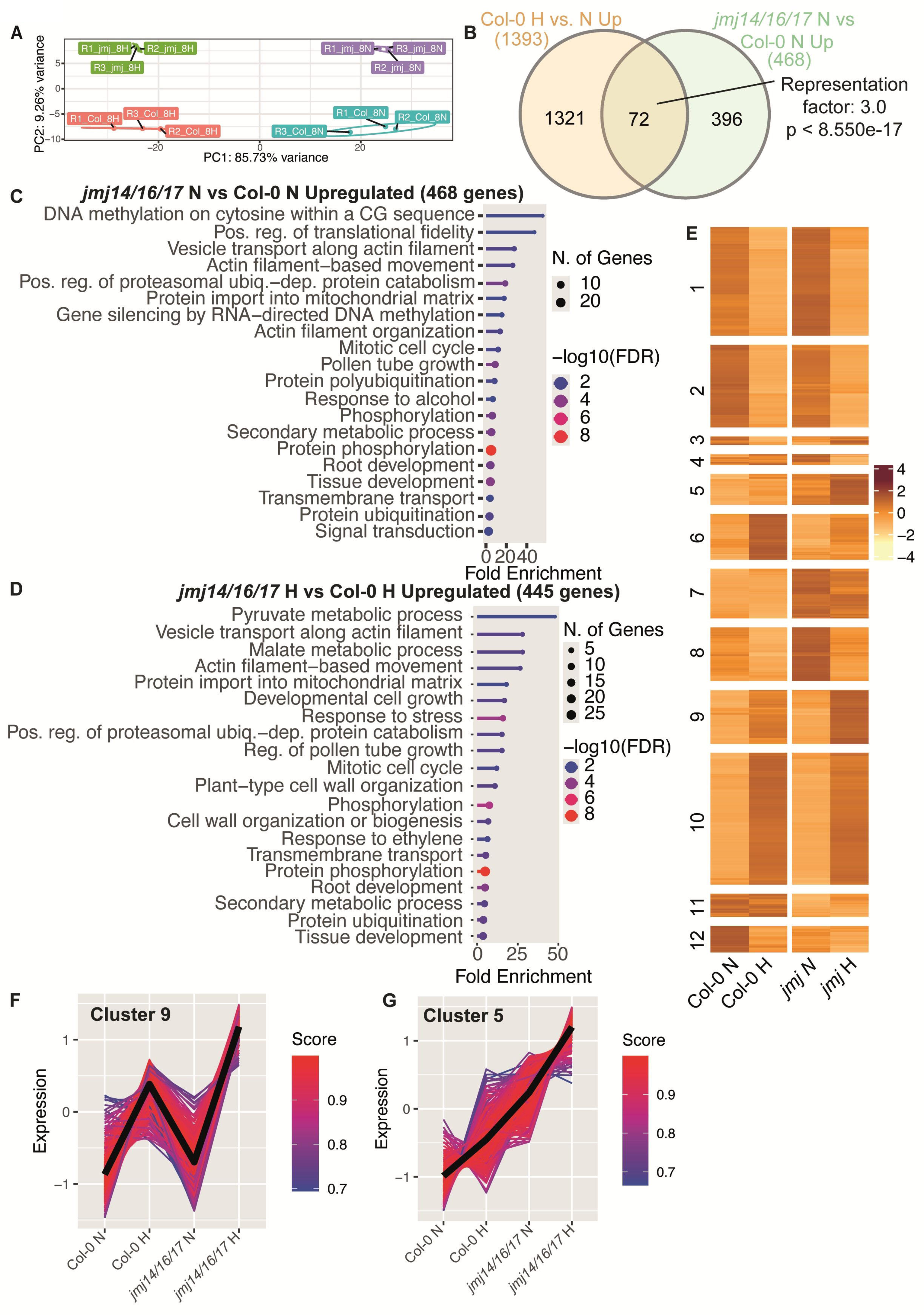
RNA-seq analyses of *jmj14-1 jmj16-1 jmj17-2* and Col-0 seedlings treated with hypoxia or normoxia. (A) Principal component analysis (PCA) showing principal component (PC) 1 versus PC2 with ellipses grouping samples by treatment (hypoxia; H and normoxia; N) and genotype (Col-0 and *jmj14-1 jmj16-1 jmj17-2*; *jmj*). (B) Venn diagram showing the overlap of differentially expressed genes (DEGs; adjusted p-value ≤ 0.05 and |log_2_(fold change)| ≥1) that were upregulated in hypoxia- versus normoxia-treated Col-0 seedlings, and DEGs that were upregulated in normoxia-treated *jmj14-1 jmj16-1 jmj17-2* versus normoxia-treated Col-0 seedlings. (C) Top 20 gene ontology (GO) terms from analyses of the 468 DEGs that were upregulated in normoxia-treated *jmj14-1 jmj16-1 jmj17-2* versus normoxia-treated Col-0 seedlings. (D) Top 20 GO terms from analyses of the 445 DEGs that were upregulated in hypoxia-treated *jmj14-1jmj16-1 jmj17-2* versus hypoxia-treated Col-0 seedlings. (E) Heatmap showing results of k-means clustering (k=12) of 6405 DEGs (see Materials and Methods for selection criteria). (F) Plot of centroid expression (bold black line) and individual gene expression (coloured lines) for genes in cluster 9. (G) Plot of centroid expression (bold black line) and individual gene expression (coloured lines) for genes in cluster 5. The colour of individual lines indicates their correlation score of gene expression relative to the centroid in plots (F-G).

We identified 1393 genes upregulated and 1519 downregulated genes between Col-0 seedlings that were treated with hypoxia compared to those kept in normoxia (Supplementary Table 4) (*p.adj* value ≤ 0.05 and |log2(fold-change)| ≥ 1). We also identified 1312 genes upregulated and 1367 downregulated genes between *jmj14-1 jmj16-1 jmj17-2* seedlings that were treated with hypoxia compared to those maintained in normoxia (*p.adj* value ≤ 0.05 and |log_2_(fold change)| ≥ 1; Supplementary Table 4). We next compared our RNA-seq data to previously published transcriptomic datasets that used similar hypoxia treatments to our study. Lee and Bailey-Serres (2019) performed polyadenylated RNA-seq on 7 d-old Col-0 seedlings that were grown in long-day conditions (16 h light: 8 h dark) and treated with hypoxia (<2% oxygen) for 9 h (Lee and Bailey-Serres, 2019). We compared the up and downregulated genes identified in Col-0 seedlings after hypoxia treatment in our study with the hypoxia induced DEGs identified by Lee and Bailey-Serres (2019) and found significant overlaps between the two groups (*p*<1.167e-248 and *p*<2.203e-159, respectively; Fig. S3A-B, Supplementary Table 10). We also observed a significant overlap between the up- and downregulated DEGs in response to hypoxia in *jmj14-1 jmj16-1 jmj17-2* seedlings and the DEGs identified by Lee and Bailey-Serres (2019) (*p*<6.923e-275 and *p*<3.426e-155, respectively; Fig. S3A-B, Supplementary Table 10).

Next, we sought to compare our RNA-seq data from *jmj14-1 jmj16-1 jmj17-2* seedlings to previously published transcriptomic datasets from individual histone demethylase mutants. We found an enrichment of DEGs identified in *jmj14-1 jmj16-1 jmj17-2* seedlings compared to wild-type seedlings in previously published datasets from *jmj14-1* seedlings (Ning *et al*., 2015) and *jmj17-1* seedlings (Huang *et al*., 2019), with the largest overlap identified between our data and *jmj14-1* seedlings (Fig. S3C-F). Unsurprisingly, we did not find any notable overlap between our data and DEGs that were identified in *jmj16-1* rosette leaves of 5-week-old plants (Liu *et al*., 2019), most likely because of the different developmental time points used in both studies (Fig. S3C-F). This suggests that perturbation of JMJ14 activity causes a larger shift in gene expression in the *jmj14-1 jmj16-1 jmj17-2* seedlings than reduction of JMJ16 or JMJ17 activity.

We compared the 3736 genes that were previously found to associate with JMJ14 binding in their genomic vicinities in flower buds by ChIP-seq (Wang *et al*., 2023) to the genes we identified as upregulated in normoxia-treated *jmj14-1/16-1/17-2* vs Col-0 seedlings (Fig. S3G). As expected, we found that genes associated with JMJ14 binding were enriched amongst the genes that were upregulated in *jmj14-1 jmj16-1 jmj17-2* seedlings in our study (representation factor: 4.2; *p*<3.161e-114; Fig. S3G). We also performed a GO analysis of 3736 JMJ14-bound genes and saw a 2.85-fold enrichment for the GO term “Cellular response to hypoxia” (FDR = 2.42e-7, Supplementary Table 11) suggesting that under normal oxygen levels, JMJ14 binds in genomic regions containing hypoxia response genes.

We next examined the DEGs identified in *jmj14-1 jmj16-1 jmj17-2* mutant seedlings to determine whether they exhibit signatures of hypoxia response. We focused our analyses on up-regulated genes as all of the differentially expressed core HRGs (Mustroph *et al*., 2009) were elevated in their expression in hypoxia treated wild-type seedlings (Supplementary Table 5). Of the 468 genes upregulated in *jmj14-1 jmj16-1 jmj17-2* triple mutant relative to Col-0 in normoxic conditions, hypoxia induced genes were 3 times overrepresented (*p*<8.55e-17) (Fig. 2B). We performed a gene ontology (GO) analysis on these 468 up-regulated genes and identified enriched terms associated with DNA methylation, ubiquitin-dependent proteosomal degradation and pollen-tube growth (Fig. 2C). A total of 518 genes were differentially expressed between Col-0 and *jmj14-1 jmj16-1 jmj17-2* seedlings that were both exposed to 8 h hypoxia treatment, 445 of which were upregulated (Supplementary Table 3, 7). GO analyses of these upregulated 445 DEGs identified terms relating to metabolic processes, actin filament-based vesicle movement and both protein import and turnover (Fig. 2D). This data shows that *jmj14-1 jmj16-1 jmj17-2* seedlings show differences in the expression of genes involved in metabolic processes and in protein post-translational regulation when responding to hypoxic stress. Notably, DEGs associated with GO terms such as ‘pyruvate metabolic process’ or ‘response to ethylene’ were over-represented in this dataset, which are particularly relevant to plant responses to hypoxia (Zabalza *et al*., 2009; Hartman *et al*., 2021).

In order to identify distinct global transcriptional signatures of hypoxia responses we performed *k*-means clustering analysis. This analysis identified 12 clusters of DEGs with distinct expression patterns between Col-0 and *jmj14-1 jmj16-1 jmj17-2* samples (Fig. 2E; Supplementary Table 5). Three clusters (cluster 1, 2, 10) were identified that showed similar transcriptional responses to hypoxia in both Col-0 and *jmj14-1 jmj16-1 jmj17-2* seedlings (Fig S4). Cluster 10 was induced in response to hypoxia and was also the largest cluster identified with 1248 DEGs, including the majority of previously identified core hypoxia response genes (42/49) such as *ALCOHOL DEHYDROGENASE 1* (*ADH1*), *HEMOGLOBIN 1* (*HB1*) and *HYPOXIA RESPONSE ATTENUATOR 1* (*HRA1*) (Fig. S4E-F, Supplementary Table 5).

Two clusters (cluster 5 and 9) contained DEGs that were upregulated in response to hypoxia in both genotypes and showed higher expression in *jmj14-1 jmj16-1 jmj17-2* hypoxia treated seedlings compared to hypoxia treated Col-0 (Fig. 2F-G). Cluster 5 also showed a higher centroid expression in *jmj14-1 jmj16-1 jmj17-2* normoxia treated seedlings compared to Col-0 normoxia treated seedlings (Fig. 2G). GO analysis of the 508 DEGs in cluster 9 revealed terms associated with hypoxia and defence responses, as well as hormone-mediated signalling pathways (Fig. S5A). Cluster 9 also included 6 of the 49 core hypoxia response genes, including *PCO1*, *JASMONATE-ZIM-DOMAIN PROTEIN 3* (*JAZ3*) and *RGA TARGET 1* (*RGAT1*) (Supplementary Table 5). Similarly, GO terms relating to defence responses were identified from the 293 DEGs in cluster 5, while GO terms relating to transport of toxic metal ions, responses to oxygen-containing compounds and protein import into mitochondrial matrix were also identified from this cluster (Fig. S5B). Two other clusters (cluster 7 and 8) showed higher centroid expression in normoxic *jmj14-1 jmj16-1 jmj17-2* samples compared to Col-0 in normoxia, however, the DEGs in these clusters were mostly downregulated in response to hypoxia (Fig. S6A, C). The expression of these genes remained higher in *jmj14-1 jmj16-1 jmj17-2* samples compared to Col-0 in hypoxic conditions. Genes identified in clusters 7 and 8 were enriched for GO terms relating to DNA methylation, root and pollen development (Fig. S6B, D).

A number of clusters (cluster 6, 11, 12) were identified which showed lower centroid expression in *jmj14-1 jmj16-1 jmj17-2* samples compared to Col-0 (Fig. S7A, C, E). JMJ14, JMJ16 and JMJ17 are known to target H3K4me3 for removal, leading to repression of target gene expression (Lu *et al*., 2010; Huang *et al*., 2019; Liu *et al*., 2019). We would therefore expect that direct targets of JMJ14, JMJ16 and JMJ17 would show enhanced gene expression in *jmj14-1 jmj16-1 jmj17-2* seedlings, suggesting that the genes in clusters 6, 11 and 12 may be indirect targets of JMJ14, JMJ16 and JMJ17. GO analysis of these clusters included terms relating to defence signalling, sucrose transport and secondary metabolite biosynthesis (Fig. S7B, D, F).

The two smallest clusters (cluster 3, 4) contained genes with opposite hypoxia mediated transcriptional responses between *jmj14-1 jmj16-1 jmj17-2* and Col-0 seedlings (Fig. S8). DEGs that were identified in cluster 3 were downregulated in Col-0 seedlings treated with hypoxia compared to normoxia (Fig. S8A). However, these same genes were generally upregulated in *jmj14-1 jmj16-1 jmj17-2* seedlings treated with hypoxia compared to normoxia (Fig. S8A). The DEGs identified in cluster 4 showed a slight increase in centroid expression in Col-0 seedlings treated with hypoxia compared to normoxia, while these same genes were consistently downregulated in *jmj14-1 jmj16-1 jmj17-2* seedlings upon hypoxia treatment (Fig. S8B). Together, these data show unique gene expression patterns in *jmj14-1 jmj16-1 jmj17-2* seedlings compared to Col-0 in response to hypoxia.

### Overview of hypoxia induced transcriptional signatures across multiple mutants

We wished to understand how the transcriptomic signatures of hypoxia treated *jmj14-1 jmj16-1 jmj17-2* seedlings compare with seedlings from mutant lines in which the transcriptional hypoxia response programme is perturbed. To do this, we performed RNA-seq on 10 d-old *ate1-2 ate2-1* (this mutant exhibits constitutive hypoxia response due to the accumulation of the ERF-VIIs in normoxic conditions) and *erf-vii* whole seedlings that were grown and subjected to the same hypoxia treatments as the *jmj14-1 jmj16-1 jmj17-2* and Col-0 seedlings. A PCA showed that the samples could be separated based on treatment (PC1: 72.84% variance) (Fig. 3A). As expected, based on the constitutive hypoxia response of *ate1-2 ate2-1* mutants, we observed a higher number of genes differentially expressed between *ate1-2 ate2-1* and Col-0 seedlings in normoxia (481 genes; Fig. S9A, Supplementary Table 4) compared to seedlings that were treated with hypoxia (249 genes; Fig. S9B, Supplementary Table 4). The opposite trend was observed with *erf-vii* seedlings, with just 60 DEGs between *erf-vii* and Col-0 seedlings in normoxia (Fig. S9A, Supplementary Table 4) and 1153 between the two genotypes when they were exposed to hypoxia (Fig. S9B, Supplementary Table 4).

**Figure 3.**
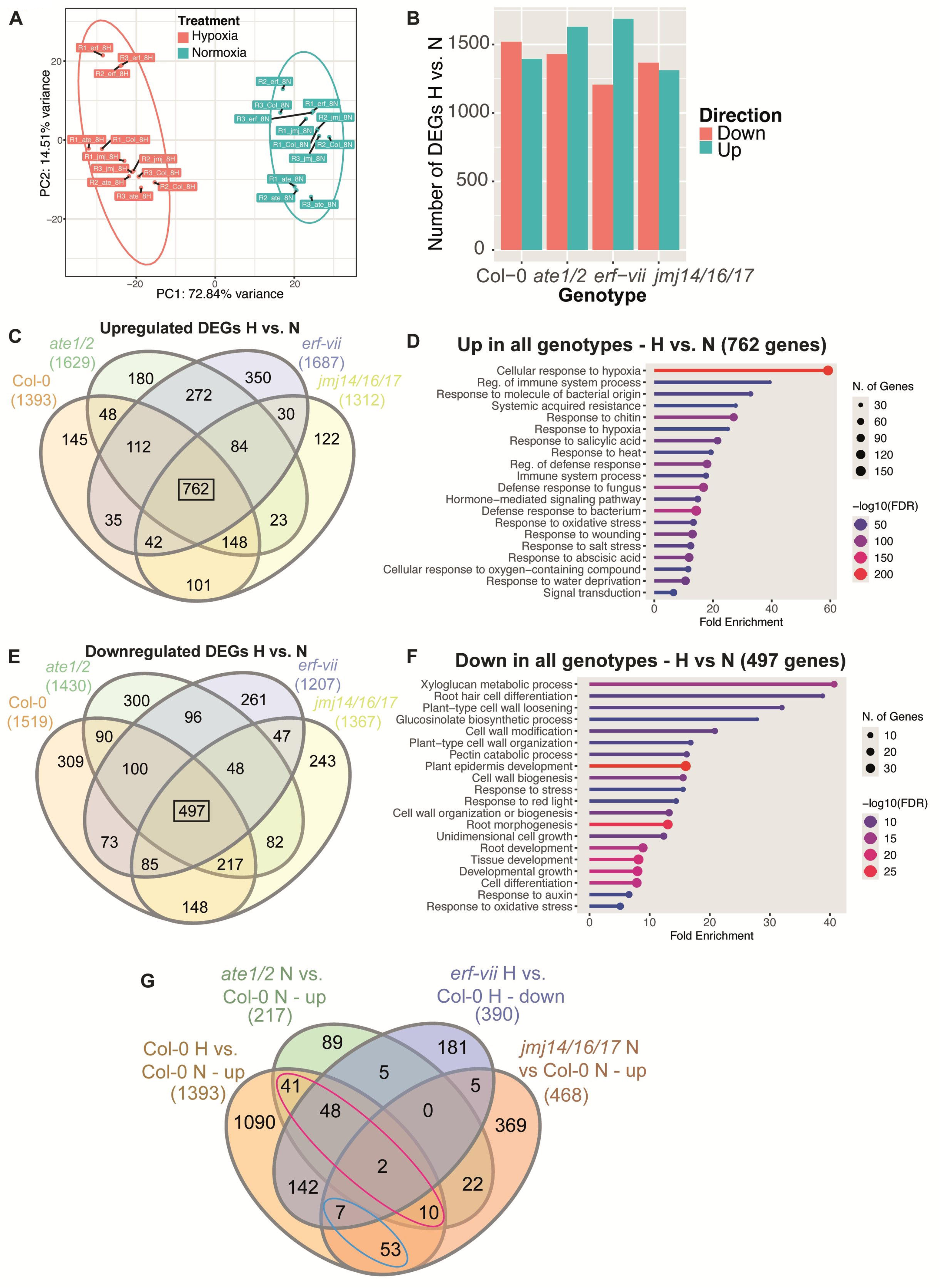
Analyses of transcriptomic response to hypoxia across seedlings with perturbed H3K4me3 histone demethylase function or defects in the canonical hypoxia response pathway. (A) PCA showing PC1 versus PC2 with ellipses grouping samples by treatment (hypoxia; H and normoxia; N). Individual samples are labelled with treatment and genotype. (B) Plot showing the number of DEGs identified when comparing hypoxia-treated to normoxia treated seedlings from different genotypes. (C) Venn diagram showing the overlap of DEGs that were upregulated in hypoxia- versus normoxia-treated seedlings in each genotype. (D) Top 20 GO terms from analyses of the 762 DEGs that were upregulated in all genotypes in response to hypoxia. (E) Venn diagram showing the overlap of DEGs that were downregulated in hypoxia- versus normoxia-treated seedlings in each genotype. (F) Top 20 GO terms from analyses of the 497 DEGs that were downregulated in all genotypes in response to hypoxia. (G) Venn diagram showing the overlap between DEG sets that are upregulated in Col-0 in response to hypoxia, upregulated in normoxia-treated *ate1-2 ate2-1* compared to normoxia-treated Col-0, downregulated in in hypoxia-treated *erf-vii* compared to hypoxia-treated Col-0 and upregulated in normoxia-treated *jmj14/16/17* compared to normoxia-treated Col-0. The genotypes used in transcriptomic analyses include Col-0, *jmj14-1 jmj16-1 jmj17-2* (*jmj or jmj14/16/17*)*, ate1-2 ate2-1* (*ate* or *ate1/2*) and *erf-vii* (*erf*).

An average of ∼2900 genes were differentially expressed between hypoxia and normoxia treated seedlings in each genotype with similar numbers of up/down regulated DEGs in each comparison (Fig. 3B, Supplementary Table 4). A total of 2454 DEGs were upregulated in response to hypoxia across all genotypes (Fig. 3C). A set of 762 genes were upregulated in response to hypoxia in every genotype analysed (Fig. 3C). A GO analyses of these 762 DEGs revealed terms relating to hypoxia and defence responses (Fig. 3D). The genes within this cohort included 37 of the 49 core HRGs (Supplementary Table 5). Of the core HRGs that did not reach the thresholds we set for upregulation (p-value ≤ 0.05 and log_2_(fold change)≥1) in all genotypes in response to hypoxia, 2 were upregulated in the *jmj14-1 jmj16-1 jmj17-2* and Col-0 seedlings only (*JAZ3* and *ATYPICAL CYS HIS RICH THIOREDOXIN 5* (*ACHT5*), while one was only upregulated in response to hypoxia in *jmj14-1 jmj16-1 jmj17-2* seedlings (*URIDINE KINASE-LIKE 3; UCK3*) (Supplementary Table 5). A set of 2596 genes were downregulated in hypoxia treated seedlings across all genotypes analysed, with a subset of 497 DEGs downregulated in response to hypoxia in every genotype analysed (Fig. 3E). GO analyses of these 497 DEGs included terms relating to cell wall organisation and root development (Fig. 3F).

We next sought to describe what proportion of hypoxia induced gene expression can be explained by the N-degron pathway or the histone demethylases JMJ14/16/17. To this end, we focused on genes that could be regulated by ERF-VIIs, which includes (i) genes that were upregulated in response to hypoxia in Col-0, (ii) genes that were upregulated in *ate1-2 ate2-1* and *jmj14-1 jmj16-1 jmj17-2* seedlings in normoxia compared to wild-type seedlings in normoxia and (iii) genes that were downregulated in *erf-vii* seedlings treated with hypoxia compared to wild-type seedlings treated with hypoxia. We found that *ate1-2 ate2-1* mutant seedlings in normoxia showed upregulation of only 101 of the 1393 genes (7%) that were induced by hypoxia treatment in wild-type seedlings (pink ellipse, Fig. 3G), with 12 of these genes also upregulated in *jmj14-1 jmj16-1 jmj17-2* seedlings in normoxia. A set of 60 genes that were induced by hypoxia in wild-type seedlings were upregulated in *jmj14-1 jmj16-1 jmj17-2* seedlings in normoxia, but not *ate1-2 ate2-1* seedlings in normoxia (blue ellipse, Fig. 3G, Supplementary Table 12). Notably, the majority of these genes (53) were not down-regulated in hypoxia-treated *erf-vii* compared to hypoxia-treated wild-type seedlings, suggesting that their regulation by hypoxia may be ERF-VII independent (Fig. 3G). According to previously published ChIP-seq data, a subset of 25 of these 53 genes were bound by JMJ14 (Wang *et al*., 2023), while 9 were bound by HYPOXIA RESPONSIVE ERF 2 (HRE2), one of the 5 ERF-VII transcription factors (Lee and Bailey-Serres, 2019), with 4 genes bound by both HRE2 and JMJ14 (Supplementary Table 12). Among the genes that were direct targets of JMJ14 but not of HRE2, we found genes encoding proteins involved in ATP hydrolysis, including CELL-DIVISION-CYCLE PROTEIN 48 B (CDC48B) and AT3G28580 (Baruah *et al*., 2009; Niehl *et al*., 2012; Yang *et al*., 2022). This data suggests that a subset of hypoxia-induced genes could be controlled by the activity of JMJ14/16/17 independently of the ERF-VII transcription factors.

### Analyses of gene expression in subset of core HRGs

We wished to describe in detail the specific effect of perturbation of JMJ14/16/17 on the core hypoxia response genes. As mentioned previously, we identified a cluster of DEGs that were upregulated in response to hypoxia and generally showed higher average expression in *jmj14-1 jmj16-1 jmj17-2* compared to Col-0 seedlings (cluster 9; Fig. 2F). This cluster contained 6 of the 49 core HRGs, including *JAZ3*, *PCO1*, *CALMODULIN-LIKE 38* (*CML38*), *FAR-RED-ELONGATED HYPOCOTYL1-LIKE*

(*FHL*), *UCK1* and *RGAT1*. In order to understand how gene expression of the core hypoxia response genes change across the mutant genotypes, we analysed the normalised RNA-seq read count data from these lines (Fig. 4, Fig. S10). We identified three core HRGs, *JAZ3*, *PCO1* and *CML38*, that were expressed at higher levels in *jmj14-1 jmj16-1 jmj17-2* seedlings treated with hypoxia compared with Col-0 seedlings treated with hypoxia (Fig. 4A-C, Supplementary Table 9). The hypoxia induced expression of these three genes was attenuated in *erf-vii* seedlings compared to wild-type seedlings (Fig. 4A-C, Supplementary Table 9). *CML38* exhibited the most striking increase in core HRG expression in the *jmj14-1 jmj16-1 jmj17-2* seedlings treated with hypoxia, with 1.64 and 1.42-fold increase compared to Col-0 and *ate1-2 ate2-1* seedlings, respectively, also treated with hypoxia (*p_adj_*=0.0004 and *p_adj_* =0.0065; Supplementary Table 9) (Fig. 4C).

**Figure 4.**
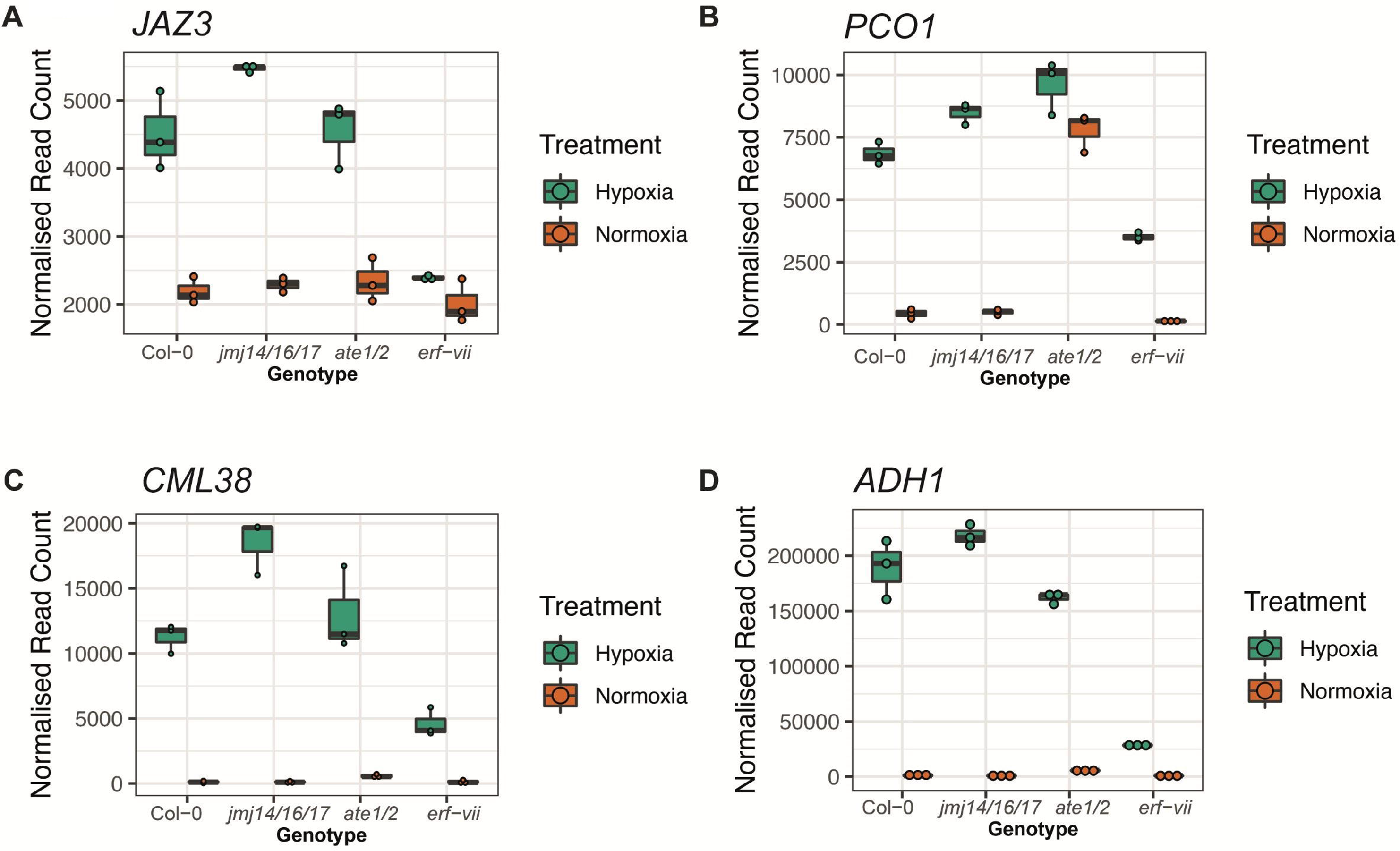
Expression of selected core hypoxia response genes in across multiple genotypes treated with hypoxia or normoxia. Boxplots of normalised read counts from RNA-seq analyses showing gene expression levels for (A) *JAZ3*, (B) *PCO1*, (C) *CML38* and (D) *ADH1*. The genotypes used in transcriptomic analyses include Col-0, *jmj14-1 jmj16-1 jmj17-2* (*jmj14/16/17*)*, ate1-2 ate2-1* (*ate1/2*) and *erf-vii*.

For the other 3 core HRGs in cluster 9, *FHL*, *UCK1* and *RGAT1*, the increase of gene expression in *jmj14-1 jmj16-1 jmj17-2* seedlings treated with hypoxia compared to Col-0 was less clear (Fig. S10 A-C, Supplementary Table 9). Gene expression of *FHL* tended to be higher in hypoxia-treated *jmj14-1 jmj16-1 jmj17-2* seedlings compared to Col-0 seedlings treated with hypoxia, although its expression in *jmj14-1 jmj16-1 jmj17-2* seedlings was variable (*p_adj_*=0.882, Fig. S10A, Supplementary Table 9). Notably, gene expression of *FHL* showed a consistent increase in hypoxia-treated *jmj14-1 jmj16-1 jmj17-2* seedlings compared to hypoxia-treated seedlings treated *ate1-2 ate2-1* seedlings (p=0.019, Fig. S10A, Supplementary Table 9). Similar patterns of gene expression were observed for *UCK1* and *RGAT1*, so that *FHL*, *UCK1* and *RGAT* showed a mild hypoxia-dependent hyper-induction in *jmj14-1 jmj16-1 jmj17-2* seedlings compared to Col-0 seedlings, while in *ate1-2 ate2-1* mutants, their hypoxia-dependent induction was dampened compared to Col-0 (Fig. S10A-C, Supplementary Table 9).

We also analysed the gene expression of a selection of core hypoxia response genes from cluster 10 (Fig. S4E) across our mutant lines, including *ADH1*, *HRA1*, *HB1* and *PCO2* (Fig. 4D, S10D-F). As expected, based on the average behaviour of genes in cluster 10, we observed similar levels of gene expression for these genes between Col-0 and *jmj14-1 jmj16-1 jmj17-2* treated seedlings (Fig. 4D, S10D-F, Supplementary Table 9). We also validated the results from our RNA-seq with RT-qPCR for *ADH1* and *HB1* and found similar patterns of gene expression across all samples (Fig. S10G-H).

### Transcriptional changes of genes coding for proteins involved in the removal or deposition of H3K4 methylation

Considering the global changes in H3K4me3 upon hypoxia treatment (Fig. 1), we assessed if hypoxia treatment influenced the expression of genes coding for proteins involved in the removal or deposition of methylation of H3K4. As expected, the levels of gene expression of *JMJ14*, *JMJ16* and *JMJ17* were lower in *jmj14-1 jmj16-1 jmj17-2* seedlings than in any of the other genotypes (Fig. 5A-C, Supplementary Table 9). As expected from publicly available gene expression data (Figure S2), we detected very low levels of *JMJ15* compared to the other H3K4 histone demethylases (Fig. 5D). Treatment status affected gene expression levels for both *JMJ16* (*p_adj_*=7.7e-06) and *JMJ17* (*p_adj_*=3.82e-07), but not *JMJ14* (*p_adj_*=0.329) with elevated levels of gene expression observed in hypoxia treated seedlings (Fig. 5A-C, Supplementary Table 9). This could suggest a feedback loop between their expression levels and decreased protein activity. Gene expression levels of *JMJ18* tended to show more variability in hypoxia treated seedlings than normoxia treated seedlings, making it difficult to draw conclusions (Fig. 5E).

**Figure 5.**
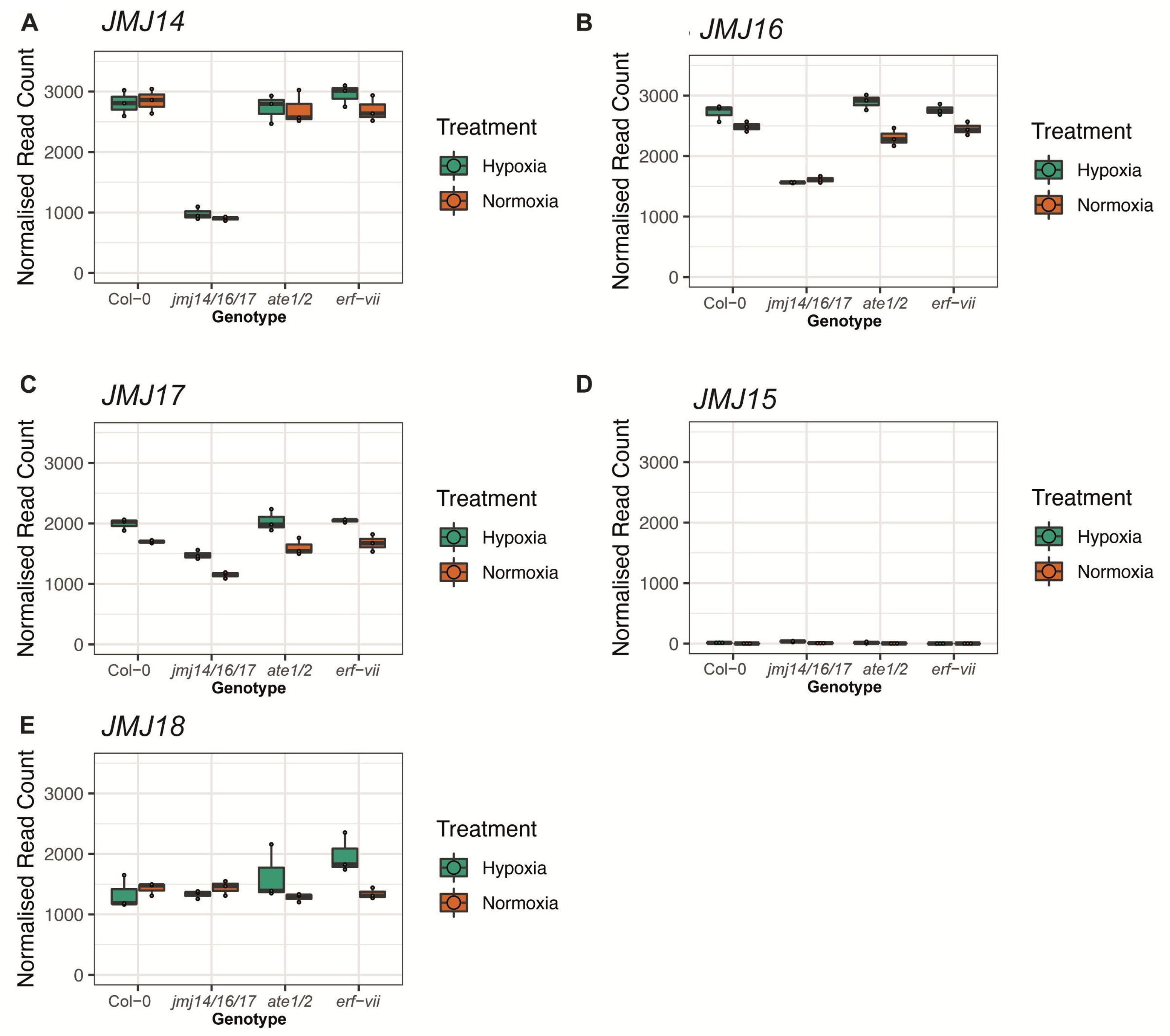
Expression of genes coding for JmjC H3K4 histone demethylases across multiple genotypes treated with hypoxia or normoxia. Boxplots of normalised read counts from RNA-seq analyses showing gene expression levels for (A) *JMJ14*, (B) *JMJ16*, (C) *JMJ17*, (D) *JMJ15* and (E) *JMJ18*. The genotypes used in transcriptomic analyses include Col-0, *jmj14-1 jmj16-1 jmj17-2* (*jmj14/16/17*)*, ate1-2 ate2-1* (*ate1/2*) and *erf-vii*.

We next analysed the expression of a number of genes coding for proteins involved in the methylation of H3K4, including *ARABIDOPSIS TRITHORAX* 3 (*ATX3*), *ATX4*, *ATX5* and *SET DOMAIN GROUP 2* (*SDG2* / *ATXR3*) (Guo *et al*., 2010; Chen *et al*., 2017). *ATX4* and *ATX5* were downregulated by up to 40% in all genotypes comparing hypoxia to normoxia treated seedlings (p*_adj_*<0.05, Fig. S11A-B, Supplementary Table 9). No consistent changes in gene expression were observed for *ATX3* or *SDG2* in most genotypes, however, a mild reduction in hypoxia *versus* normoxia treated seedlings was observed for *SDG2* in Col-0 (*p_adj_*=0.089, Fig. S11C-D, Supplementary Table 9). This data supports the hypothesis that hypoxia-induced H3K4me3 accumulation is due to an inhibition of histone demethylase activity rather than an increase in histone methyltransferase activity, however, protein and kinetic based evidence would also be necessary to fully understand the contribution of histone methyltransferases in this process.

### Transcriptional changes of genes coding for proteins involved in the removal or deposition of H3K27 methylation

We next analysed the transcriptional changes for genes coding for proteins involved in the (de)methylation of H3K27me3, a histone mark associated with repression of gene expression that acts antagonistically to H3K4me3. Methylation at H3K27 can be removed by a subset of Jmj-C domain histone demethylases, including RELATIVE OF EARLY FLOWERING 6 (REF6), EARLY FLOWERING 6 (ELF6) and JMJ13, and which are also likely to be oxygen dependent. REF6 and ELF6 expression tended to be lower in hypoxia treated seedlings compared to normoxia (*p_adj_*= 0.457, *p_adj_*= 0.173 in Col-0, Fig. S12A-B, Supplementary Table 9). An exception to this was *ELF6* expression which was induced by hypoxia in *ate1-2 ate2-1* seedlings (*p_adj_*= 0.017, Fig. S12B, Supplementary Table 9) and unchanged between hypoxia and normoxia in *erf-vii* seedlings (*p_adj_*= 1, Fig. S12B, Supplementary Table 9). In contrast to *REF6* and *ELF6*, hypoxia treatment induced *JMJ13* expression (Figure S12C), with Col-0 showing 2.2-fold higher expression of *JMJ13* compared to normoxia treated seedlings (*p_adj_*=0.00099, Supplementary Table 9).

The genes encoding for the two histone methyltransferases that confer methylation of H3K27me3, CURLY LEAF (CLF) and SWINGER (SWN), both showed differences in gene expression that were dependent on hypoxia treatment, albeit with opposite directions (Fig. S13A-B). *CLF* expression was ∼30% lower in hypoxia treated Col-0 seedlings compared to normoxia treated seedlings (*p_adj_*=0.01, Fig. S13A, Supplementary Table 9), while *SWN* expression was ∼30% higher in hypoxia treated Col-0 seedlings compared to normoxia treated seedlings (*p_adj_*=6.4e-05, Fig. S13B, Supplementary Table 9). We also observed an influence of the interaction between genotype and treatment on the gene expression of *SWN* (*p_adj_*= 0.027, Supplementary Table 9), with *erf-vii* showing a lower magnitude of hypoxia-induced *SWN* expression compared to Col-0, for example (Fig. S13B, Supplementary Table 9). No clear differences in gene expression were observed for *FERTILISATION INDEPENDENT ENDOSPERM* (*FIE*) or *MULTICOPY SUPRESSOR OF IRA1* (*MSI1*), two genes encoding for core subunits of PRC2 (Fig. S13C-D, Supplementary Table 9) while treatment status was found to influence gene expression levels for *EMBRYONIC FLOWER 2* (*EMF2*) (p_adj_=0.008, Fig. S13E, Supplementary Table 9) and for *VRN2* (p_adj_= 0.01, Fig. S13F, Supplementary Table 9).

## Discussion

Plants use oxygen sensing mechanisms to regulate stress survival responses in the case of acute environmental hypoxia, or developmental pathways in the context of tissue specific hypoxic niches (Loreti and Perata, 2020). The hypoxia-induced expression of a core set of genes and their orthologues that are central to the hypoxia response pathway has been documented across multiple plant species (Mustroph *et al*., 2009, 2010; Safavi-Rizi *et al*., 2020; Ambros *et al*., 2022; Miricescu *et al*., 2023). In mammals, the inhibition of histone demethylases by low oxygen conditions has been shown to play a key role in the induction of certain hypoxia-response genes (Batie *et al*., 2019; Chakraborty *et al*., 2019), however, little is known about the contribution of orthologous histone demethylases to hypoxia responses in plants. Here, we investigated how global levels of H3K4me3, a histone mark associated with activation of gene expression, respond to hypoxia (Fig. 1, Fig. S1). We also used transcriptomic approaches to explore a role for H3K4me3 histone demethylases in regulating HRGs (Fig. 2-5; Fig. S2-13).

### An N-degron independent pathway controls a portion of hypoxia responses in plants

The observation that 93% of genes that are upregulated in response to hypoxia treatment in wild-type seedlings are not upregulated in *ate1-2 ate2-1* seedlings under normoxia (Fig. 3G) suggests that an alternative pathway to the N-degron pathway is involved in mediating the activation of HRGs in *A. thaliana*. Similar to previous observations (Gibbs *et al*., 2011), ∼55% core HRGs were not induced in *ate1-2 ate2-1* seedlings under normoxia compared to Col-0 seedlings in normoxia, including *JAZ3*, *CML38*, *FHL*, *UCK1*, *RGAT1*, *HRA1* and *HB1* (Fig. 4A, 4C, S10A-E, Supplementary Table 7, Supplementary Table 9). A subset of these genes (*CML38*, *HRA1* and *HB1*) were up-regulated in hypoxia treated *erf-vii* mutants, albeit to lower levels than in hypoxia treated Col-0 seedlings (Fig. 4C, S10D-E, Supplementary Table 9). Together, these data suggest that a pathway independent of N-degron mediated protein degradation of ERF-VIIs and VRN2 exists that can regulate the expression of hypoxia response genes.

### Global H3K4me3 levels are correlated with the activation of hypoxia responses

ChIP-seq analyses of H3K4me3 showed a mild increase in H3K4me3 enrichment across the gene body of core HRGs after 2 h hypoxia treatment of *A. thaliana* seedlings, while global analysis of all gene bodies showed a decrease in H3K4me3 enrichment in seedlings after 2 h hypoxia (Lee and Bailey-Serres, 2019). This suggests that H3K4me3 may play a specific role in the regulation of HRGs. In agreement with this idea, Western blot analysis showed global changes in H3K4me3 levels in wild-type Col-0 seedlings after 8 h of hypoxia treatment (Fig. 1B-C). We also observed that mean global levels of H3K4me3 were approximately 1.6-fold higher *ate1-2 ate2-1* seedlings, in which the hypoxia response pathway is constitutively active, treated with hypoxia *versus* hypoxic wild-type seedlings (Fig. 1D, Supplementary Table 2). Our data indicates that the activation of the hypoxia response pathway is positively correlated with global H3K4me3 accumulation.

### The expression of genes coding for epigenetic enzymes shows dynamic changes in response to hypoxia

It has previously been observed in mammalian cells that the mRNA and protein levels of Jmj-C histone demethylases are enriched after hypoxia treatment (Sar *et al*., 2009; Yang *et al*., 2009; Ponnaluri *et al*., 2011). We observed a similar hypoxia-induced elevation of gene expression for *JMJ16* (*p_adj_*=7.7e-06), *JMJ17* (*p_adj_*=3.82e-07), and *JMJ13* (*p_adj_*= 2.62e-09), but not for *JMJ14* (Fig. 5A-C, Fig S12C, Supplementary Table 9). For two histone demethylases that target H3K27me3, *REF6* and *ELF6*, hypoxia treatment lowered their gene expression in hypoxia treated seedlings (Figure S12A-B, Supplementary Table 9). Notably, a previously published ChIP-seq dataset for HRE2 (Lee and Bailey-Serres, 2019), one of the five ERF-VII transcription factors, showed enrichment for HRE2 at the promoter/first exon of the genes coding for *JMJ13*, *JMJ14* and *REF6* (Supplementary Table 11), suggesting that HRE2 may regulate the expression of these target genes directly. However, in our data we did not observe reproducible differences between our *erf-vii* and wild-type seedlings for the expression of *JMJ13* (*p_adj_*=0.3818), *JMJ14* (*p_adj_*=1) or *REF6* (*p_adj_*=0.8575) under hypoxic conditions (Supplementary Table 9). It is therefore unclear what mechanism controls the expression of certain histone demethylase genes in response to hypoxia.

In mammals, both gene and protein expression levels of the H3K4me3 histone methyltransferases SETD1A and SMYD3 increase in preeclampsia hypoxic placentas that also show global accumulation of H3K4me3 (Matsui *et al*., 2021). In *A. thaliana*, SDG2 (ATXR3) is the major enzyme responsible for deposition of H3K4me3 (Guo *et al*., 2010). SET domain proteins ATX3, ATX4 and ATX5 (SDG14/16/29) are critical for plant development and function redundantly to control H3K4 methylation at thousands of loci in an independent pathway to SDG2 (Chen *et al*., 2017). Here, we observed that the gene expression of *ATX3/4/5* was down-regulated in response to hypoxia by up to 40% across multiple genotypes (Fig. S11A-C), while the levels of *SDG2* gene expression are only down-regulated in wild-type seedlings (Fig. S11D). Two SET domain proteins, CLF and SWN, are partially redundant in their deposition of H3K27me3 (Shu *et al*., 2019). Similarly to *ATX3/4/5*, expression of *CLF* is down-regulated in response to hypoxia across multiple genotypes (Fig. S13A), while gene expression of *SWN* is upregulated (Fig. S13B). The observed transcriptional regulation of histone methyltransferases makes it difficult to conclude that methyltransferase activity is increased upon hypoxia. This needs to be determined by monitoring the protein levels of methyltransferases, as well as their *in vivo* activity. These findings nevertheless suggest that the accumulation of H3K4me3 in wild-type seedlings treated with hypoxia is more likely to be due to the repression of histone demethylases.

### JMJ14/16/17 are involved in the molecular response to hypoxia

We asked whether JMJ14/16/17 could directly regulate global H3K4me3 levels in response to hypoxia. For each of the histone demethylase mutants, we compared the global levels of H3K4me3 in hypoxia *versus* normoxia using Western blotting. The *jmj16-1* mutant seedlings reproducibly behaved similarly to Col-0 and exhibited increased H3K4me3 levels in hypoxia (Fig. S1D-E, Supplementary Table 2). The hypoxia-induced enrichment of H3K4me3 was dampened in *jmj17-2* seedlings while median levels of H3K4me3 in *jmj14-1* and *jmj14-1 jmj16-1 jmj17-2* were not increased in response to hypoxia treatment (Fig. 1E-F, S1F-G, S1J, Supplementary Table 2). This suggests that JMJ14 plays a major role in the global enrichment of H3K4me3 under hypoxia treatment while JMJ16 and JMJ17 may play more minor, perhaps localised, roles, or play roles at a different developmental stages.

Previous work showed that exposure of mammalian cells to hypoxic conditions led to a global accumulation in H3K4me3 due to the inhibition of the oxygen-sensitive histone demethylase KDM5A (Batie *et al*., 2019). Transcriptomic signatures can give insights into how plants are responding to their environment. Here, we analysed the transcriptomes of *jmj14-1 jmj16-1 jmj17-2* seedlings exposed to hypoxic or normoxic conditions, and compared them to seedlings with known perturbations in their hypoxic response pathways, as well as to wild-type seedlings. We found unique transcriptomic signatures in *jmj14-1 jmj16-1 jmj17-2* seedlings compared to Col-0 seedlings (Fig. 2A, E). Notably, genes that are upregulated in response to hypoxia in Col-0 seedlings are enriched amongst genes that are upregulated under normoxia in *jmj14-1 jmj16-1 jmj17-2* seedlings compared to Col-0 seedlings (Fig. 2B). This suggests a functional overlap between these H3K4 demethylases and the hypoxia response pathway in *A. thaliana*. We identified clusters of differentially expressed genes that respond differently to hypoxia treatment in *jmj14-1 jmj16-1 jmj17-2* seedlings compared to Col-0 (Fig. 2F-G, Fig. S4-8). This included cluster 9 which contained 6 of the 49 core hypoxia response genes and showed a higher average hypoxia-induced gene expression in *jmj14-1 jmj16-1 jmj17-2* seedlings compared to Col-0 seedlings (Fig. 2F, Supplementary Table 5), as would be expected if H3K4me3 levels were increased at these genes in the triple mutant.

Notably, a subset of core hypoxia response genes was induced to higher levels in *jmj14-1 jmj16-1 jmj17-2* seedlings treated with hypoxia compared to Col-0 seedlings, including *JAZ3*, *PCO1* and *CML38* (Fig. 4A-C, Supplementary Table 9). Our analyses of previously published JMJ14 ChIP-seq data (Wang *et al*., 2023) showed that the GO term for cellular response to hypoxia is enriched in genes bound by JMJ14 (Supplementary Table 11). However, we did not observe elevated levels of gene expression in core hypoxia response genes in *jmj14-1 jmj16-1 jmj17-2* compared to Col-0 seedlings under normoxia (Supplementary Table 6). This would suggest that additional factors, for example the ERF-VII transcription factors, are required for the activation of these core hypoxia response genes in *jmj14-1 jmj16-1 jmj17-2* seedlings under normal oxygen levels. We did however find a set of 60 hypoxia-inducible genes that were upregulated in *jmj14-1 jmj16-1 jmj17-2* seedlings, but not *ate1-2 ate2-1* seedlings (blue ellipse, Fig. 3G, Supplementary Table 12). The majority of these genes (53) were not down-regulated in hypoxia-treated *erf-vii* compared to hypoxia-treated wild-type seedlings (Fig. 3G) and indeed, only 9 were found to be direct targets of HRE2 according to published ChIP-seq datasets (Lee and Bailey-Serres, 2019) (Supplementary Table 12). A subset of 25 of these genes were previously described as direct targets of JMJ14 (Wang *et al*., 2023), including genes encoding (CDC48B) and AT3G28580, two proteins involved in ATP hydrolysis (Baruah *et al*., 2009; Niehl *et al*., 2012; Yang *et al*., 2022). This data suggests that a subset of hypoxia-induced genes could be controlled by the activity of JMJ14/16/17 independently of the N-degron pathway.

## Conclusions

The global histone methylation and transcriptomic analyses data we present here together highlight epigenetic and gene expression changes that occur in response to hypoxic stress in plants. The unique transcriptomic signatures we observed in *jmj14*/16/17 seedlings also suggest histone demethylases regulate hypoxia responses in plants. The data we show indicate a potential novel role for JMJ14 as an oxygen sensor that removes H3K4me3 under normal oxygen levels while it is inhibited under hypoxia, with JMJ16/17 potentially playing more minor or stage specific roles in this process. Global levels of H3K4me3 are increased under hypoxia and this could lead to upregulation of targets genes, either in an ERF-VII dependent or independent manner (Fig. S14). The elucidation of gene-specific dynamics of H3K4me3 localisation during hypoxia treatments of histone demethylase mutants is essential to further understand the contribution of histone demethylases to the regulation of hypoxia responses. This study identifies plant H3K4 histone demethylases as novel mediators of hypoxia sensing and regulation, thus highlighting new directions for plant research.

## Supplemental Data

**Table S1.** List of primer sequences used in this study **Table S2.** Quantification of H3K4me3 Western blots **Table S3.** Normalised read counts from RNA-seq analyses.

**Table S4.** Numbers of DEGs identified in all comparisons analysed.

**Table S5.** Results of DeSeq2 analyses for hypoxia versus normoxia treatments in each genotype.

**Table S6.** Results of DeSeq2 analyses for normoxia versus normoxia treatments between all genotypes.

**Table S7.** Results of DeSeq2 analyses for hypoxia versus hypoxia treatments between all genotypes.

**Table S8.** List of comparisons used for k-means clustering.

**Table S9.** Results of statistical analyses of gene expression for selected genes.

**Table S10.** Analyses of previously published transcriptomic data-sets.

**Table S11.** Analyses of previously published ChIP-seq data-sets.

**Table S12.** Analyses of HRG subset potentially controlled by JMJ14/16/17.

## Acknowledgements

We thank Prof. Francesco Licausi, Dr. Daai Zhang and our colleagues for helpful discussions about this work.

## Author contributions

DSOM: investigation, resources, writing – review & editing; EG: conceptualisation, investigation, resources, writing – review & editing; AJB: conceptualisation, investigation, methodology, formal analysis, data curation, visualisation, funding acquisition, project administration, writing – original draft, writing – review and editing.

## Data Availability

RNA-seq data have been deposited with the Gene Expression Omnibus (GEO) repository (at http://www.ncbi.nlm.nih.gov/) under GSE299155.

## Abbreviations

A. thaliana: Arabidopsis thaliana
ERF-VII: group VII ethylene response factor
ATE: ARGININE T-RNA PROTEIN TRANSFERASE
JMJ: Jumonji-C domain
PRT6: PROTEOLYSIS 6
PCO: PLANT CYSTEINE OXIDASE
HRG: hypoxia response gene
VRN2: VERNALISATION 2 (VRN2)
PRC2: polycomb repressive complex 2 H3K: histone 3 lysine
me3: trimethylation GO: Gene ontology

---

**Fig. S1.** Dynamic changes in global H3K4me3 accumulation in response to hypoxia. Western blot analyses of H3K4me3 abundance in total protein extracts from 10 d old seedlings treated with hypoxia (0.5% oxygen; H) or normoxia (N) for 8 h is shown for Col-0 and (A) *ate1-2 ate2-1,* (B) *erf-vii*, (D) *jmj16-1*, (F) *jmj17-2*, (H) *jmj14-1* and (J) *jmj14-1 jmj16-1 jmj17-2*. Ponceau S staining of total protein is shown for all blots. (C) Quantification of H3K4me3 levels in Col-0 and *erf-vii* normalised to normoxia-treated Col-0 from all Western blots shown in Fig S1B is shown (n=3). (E) Quantification of H3K4me3 levels normalised to normoxia-treated Col-0 and *jmj16-1* from all Western blots shown in Fig S1D is shown (n=3). (G) Quantification of H3K4me3 levels normalised to normoxia-treated Col-0 and *jmj17-2* from all Western blots shown in Fig S1F is shown (n=3). (I) Quantification of H3K4me3 levels normalised to normoxia-treated Col-0 and *jmj14-1* from all Western blots shown in Fig S1H is shown (n=4).Experiments were performed in independent biological triplicate, each with two biological replicates. Blots for replicates that are not displayed in Fig. 1 are shown here. Details of which replicates are displayed in each figure can be found in Supplementary Table 2. Replicate 3.2 Col-0, *jmj14-1* and *jmj17-2* samples were run on the same gel and as such the Col-0 samples are duplicated here in panel (F) and (H) for direct comparison with both genotypes.

**Figure S2.** Gene expression of H3K4me3 histone demethylases. Gene expression atlas analyses of *JMJ14*-*JMJ19* expression analyses in (A) the whole rosette and (B) multiple tissue types from BAR eFP.

**Figure S3.** Comparison of transcriptomic analyses to previously published data-sets. Venn diagrams showing the overlap of DEGs that were (A) upregulated or (B) downregulated in hypoxia-versus normoxia-treated Col-0 and *jmj14-1 jmj16-1 jmj17-2* seedlings identified in this study and in Lee and Bailey-Serres (2019). Venn diagrams showing the overlap of DEGs that were (C) upregulated or (D) downregulated in mutant versus control Col-0 seedlings (normoxia) from *jmj14-1 jmj16-1 jmj17-2* (this study), *jmj14-1* (Ning et al. 2015), *jmj16-1* (Liu et al. 2019a) and *jmj17-2* (Huang et al. 2019a). (E) Table showing the representation factor and p-value calculated from the number of upregulated overlapping genes in *jmj14-1 jmj16-1 jmj17-2* (this study) and the data shown in (C). (F) Table showing the representation factor and p-value calculated from the number of downregulated overlapping genes in *jmj14-1 jmj16-1 jmj17-2* (this study) and the data shown in (D). (G) Venn diagrams showing the overlap of DEGs that were upregulated in normoxia-treated *jmj14-1/16-1/17-2* vs Col-0 seedlings identified in this study and JMJ14-bound genes that were identified in Wang *et al*. (2023).

**Figure S4.** Clusters of DEGs with similar gene expression patterns in *jmj14-1 jmj16-1 jmj17-2* compared to Col-0 seedlings treated with hypoxia or normoxia. (A) Plot of centroid expression (bold black line) and individual gene expression (coloured lines) for genes in cluster 1. (B) Top 20 GO terms from analyses of the 1027 genes in cluster 1. (c) Plot of centroid expression (bold black line) and individual gene expression (coloured lines) for genes in cluster 2. (d) Top 20 GO terms from analyses of the 783 genes in cluster 2. (e) Plot of centroid expression (bold black line) and individual gene expression (coloured lines) for genes in cluster 10. (f) Top 20 GO terms from analyses of the 1248 genes in cluster 10. The colour of individual lines indicates their correlation score of gene expression relative to the centroid in plots (A), (C) and (E).

**Figure S5.** Gene ontology analyses of cluster 9 and cluster 5. Top 20 GO terms from analyses of the (A) 508 genes in cluster 9 and (B) 293 genes in cluster 5. The average centroid expression of clusters 9 and 5 are shown in Fig. 2F-G.

**Figure S6:** Clusters of DEGs with higher average gene expression in *jmj14-1 jmj16-1 jmj17-2* compared to Col-0 seedlings treated with hypoxia or normoxia. (A) Plot of centroid expression (bold black line) and individual gene expression (coloured lines) for genes in cluster 7. (B) Top 20 GO terms from analyses of the 470 genes in cluster 7. (C) Plot of centroid expression (bold black line) and individual gene expression (coloured lines) for genes in cluster 8. (D) Top 20 GO terms from analyses of the 508 genes in cluster 8. The colour of individual lines indicates their correlation score of gene expression relative to the centroid in plots (A) and (C).

**Figure S7.** Clusters of DEGs with lower average gene expression in *jmj14-1 jmj16-1 jmj17-2* compared to Col-0 seedlings treated with hypoxia or normoxia. (A) Plot of centroid expression (bold black line) and individual gene expression (coloured lines) for genes in cluster 6. (B) Top 20 GO terms from analyses of the 433 genes in cluster 6. (C) Plot of centroid expression (bold black line) and individual gene expression (coloured lines) for genes in cluster 11. (D) Top 20 GO terms from analyses of the 221 genes in cluster 11. (E) Plot of centroid expression (bold black line) and individual gene expression (coloured lines) for genes in cluster 12. (F) Top 20 GO terms from analyses of the 250 genes in cluster 12. The colour of individual lines indicates their correlation score of gene expression relative to the centroid in plots (A), (C) and (E).

**Figure S8.** Clusters of DEGs that exhibit opposite expression patterns in response to hypoxia in *jmj14-1 jmj16-1 jmj17-2* seedlings compared to Col-0. (A) Plot of centroid expression (bold black line) and individual gene expression (coloured lines) for genes in cluster 3. (B) Plot of centroid expression (bold black line) and individual gene expression (coloured lines) for genes in cluster 4. (C) Top 20 GO terms from analyses of the 81 genes in cluster 3. (D) Top 20 GO terms from analyses of the 104 genes in cluster 4. The colour of individual lines indicates their correlation score of gene expression relative to the centroid in plots (A) and (B).

**Figure S9.** Numbers of DEGs identified when comparing within treatments. Plot showing the number of DEGs identified when comparing (A) normoxia-treated genotypes and (B) hypoxia-treated seedlings from different genotypes

**Figure S10.** Expression of selected core HRGs across multiple genotypes treated with hypoxia or normoxia. Boxplots of normalised read counts from RNA-seq analyses showing gene expression levels for (A) *FHL*, (B) *UCK1*, (C) *RGAT1*, (D) *HRA1*, (E) *HB1* and (F) *PCO2*. Boxplots of relative gene expression measured by RT-qPCR showing gene expression levels for (G) *ADH1* and (H) *HB1*. The genotypes used in gene expression analyses include Col-0, *jmj14-1 jmj16-1 jmj17-2* (*jmj14/16/17*)*, ate1-2 ate2-1* (*ate1/2*) and *erf-vii*.

**Figure S11.** Expression of genes encoding for selected SET domain proteins involved in the deposition of H3K4 methylation in across multiple genotypes treated with hypoxia or normoxia. Boxplots of normalised read counts from RNA-seq analyses showing gene expression levels for (A) *ATX4*, (B) *ATX5*, (C) *ATX3* and (D) *SDG2*. The genotypes used in transcriptomic analyses include Col-0, *jmj14-1 jmj16-1 jmj17-2* (*jmj14/16/17*)*, ate1-2 ate2-1* (*ate1/2*) and *erf-vii*.

**Figure S12.** Expression of genes encoding for JmjC H3K27 histone demethylases across multiple genotypes treated with hypoxia or normoxia. Boxplots of normalised read counts from RNA-seq analyses showing gene expression levels for (A) *REF6*, (B) *ELF6* and (C) *JMJ13*. The genotypes used in transcriptomic analyses include Col-0, *jmj14-1 jmj16-1 jmj17-2* (*jmj14/16/17*)*, ate1-2 ate2-1* (*ate1/2*) and *erf-vii*.

**Figure S13.** Expression of selected genes encoding for proteins involved in the deposition of H3K27me3 across multiple genotypes treated with hypoxia or normoxia. Boxplots of normalised read counts from RNA-seq analyses showing gene expression levels for (A) *CLF*, (B) *SWN*, (C) *FIE*, (D) *MSI1*, (E) *EMF2* and (F) *VRN2*. The genotypes used in transcriptomic analyses include Col-0, *jmj14-1 jmj16-1 jmj17-2* (*jmj14/16/17*)*, ate1-2 ate2-1* (*ate1/2*) and *erf-vii*.

**Figure S14:** Potential model for the role of JMJ14/16/17 in the regulation of plant hypoxia responses. Under normoxia, oxygen dependent histone demethylases JMJ14/16/17 remove H3K4me3 at target genes, including hypoxia response genes (HRGs), ensuring their expression remains switched off. Under hypoxic conditions the oxygen dependent histone demethylase activity of JMJ14/16/17 is reduced leading to an accumulation of H3K4me3 at target genes which potentially contributes to their activation. As described in Fig. 1, the ERF-VII transcription factors and PRC2 subunit, VRN2, accumulate in response to hypoxia and are also involved in regulating gene expression. The ERF-VIIs are involved in the activation of HRGs, while VRN2 is involved in the repression of target genes.

